# Probing efficient microbial CO_2_ utilization through metabolic and process modeling

**DOI:** 10.1101/2023.08.01.547650

**Authors:** Philip J. Gorter de Vries, Viviënne Mol, Nikolaus Sonnenschein, Torbjørn Ølshøj Jensen, Alex Toftgaard Nielsen

**Author notes:** Correspondence to: Alex Toftgaard Nielsen, The Novo Nordisk Foundation Center for Biosustainability, Technical University of Denmark, 2800 Kgs Lyngby, Denmark. These authors contributed equally and should be considered joint first authors (Philip J. Gorter de Vries, Viviënne Mol).

## Abstract

Microbial gas fermentation is proving to be a promising technology to upcycle carbon-rich waste gasses into value-added biochemicals, though production yields of varied products are currently limited. Through the holistic pairing of process modeling with host agnostic black box metabolic modeling, here we investigate an efficient thermophilic CO_2_ upcycling process, based on acetogenic carbon utilization. From a process engineering perspective, higher temperatures were found to favor overall gas transfer rates, even with lower gas solubility, particularly for the more expensive and often limiting H_2_ gas. Metabolically, for growth coupled products, thermophilic production favors higher product yields as a result of a higher maintenance energy input. A process simulation for acetate production in a large-scale bubble column reactor predicts an optimal feed gas composition of approximately 9:1 mol H_2_ to mol CO_2_ and a process with higher production yields and rates at higher temperatures. To assess the expansion of the product portfolio beyond acetate, both a product volatility analysis and a metabolic pathway model were implemented. *In-situ* recovery of volatile products is shown to be within range for acetone but challenging due to the extensive evaporation of water, while the production of more valuable compounds is energetically unfavorable compared to acetate. We discuss alternative approaches to overcome these challenges to utilize acetogenic CO_2_ fixation for the production of a wider range of carbon negative chemicals.

## 1 Introduction

To meet ambitious climate goals set by governments and institutions across the globe, the current reduction of emitted greenhouse gases (GHG) is not deemed sufficient^1^. As a result, carbon capture technologies have gained interest and are increasingly being implemented. However, the predominant carbon capture technologies rely on costly storage of sequestered carbon dioxide (CO_2_), in large underground reservoirs, where it remains unused^2^. To allow circular re-use of carbon, released or sequestered CO_2_ can be upgraded into more valuable biochemicals using Carbon Capture and Utilization (CCU). CO_2_ is a thermodynamically stable form of carbon, and as such, relies on energy input to be condensed into longer carbon chains, for example in CO_2_ electroreduction to acetate^3^ or ethanol^4^. Naturally, microbial cells function as biological catalysts capable of performing chemical reactions with lower energy inputs through their reliance on enzymes. Photosynthesis, the most abundant biologic carbon fixation pathway, harnesses light to reduce CO_2_, an energy source that is however poorly scalable in industrial-scale CCU^5^. Although less abundant, various chemoautotrophic ways of life exist, which use chemicals as energy sources. Organisms harboring such metabolism are more suitable for industrial-scale CCU and can be used to utilize carbon from gasified biomass or industrial waste gas, for example, from the steel industry^6^. A main group of interest is organisms using the Wood-Ljungdahl pathway (WLP) to reduce CO_2_ or carbon monoxide (CO) to form acetyl-CoA, with redox potential from the oxidation of CO or hydrogen (H_2_)^7–9^. The generated acetyl-CoA can be further fermented into various products, such as methane, acetate, or ethanol^10^. The class of bacteria that possess the WLP are acetogens, a polyphyletic group characterized by their main metabolic product: acetate.

The synthetic biology era has facilitated the introduction of novel phenotypes to various microbial hosts. Recent efforts have engineered synthetic microbial cell factories to introduce CO_2_ as a carbon source into model organisms such as *Escherichia coli* and *Pichia pastoris*, however, the rates that can be achieved are by far outcompeted by the natural WLP harboring organisms^11–14^. To harness the natural efficient CO_2_ fixating capabilities, various acetogens have been engineered to produce a wider range of valuable bioproducts^15–17^. Recently, the carbon-negative production of acetone and isopropanol by an engineered mesophilic *Clostridium* has been demonstrated up to pilot scale^18^. Acetogens are a diverse group showing large phenotypic variation, including their optimal growth temperature where mesophilic organisms grow in ranges up to 45 ºC, and thermophilic hosts have optimum growth temperatures above this threshold. The diversity of commonly used acetogens has been reviewed in regards to their growth temperature optima, ranging from 20-66 °C, but also other characteristics such as their prevalent metabolic substrates and products^19^. To date, most engineered acetogens are mesophilic, where only a few examples of engineered thermophilic acetogens producing alternative biochemicals exist^20^. Temperature is a factor that directly affects virtually all physicochemical aspects of a fermentation process, including the biology. Working under thermophilic conditions provide various advantages over mesophilic production conditions, including decreased contamination risks, lower cooling costs, higher mass transfer rates, a key factor considering the gaseous substrates^21^, and improved growth and production rates^22,23^. Additionally, higher fermentation temperatures allow more efficient *in situ* product recovery through gas stripping, specifically for volatile products. This facilitates simple downstream processing, whilst keeping product buildup in the fermentation broth low, in turn preventing product toxicity and feedback inhibition of production pathways^24,25^. As a result of these advantages, previously the economic feasibility of producing a volatile organic compound (VOC), acetone, from a theoretical thermophilic production strain was shown^26^. In the present study, we investigate the effect of process temperature on industrial acetogenic CO_2_ fixation by using biological and mass transfer models, focusing especially on the effects of temperature on metabolism, gas uptake rates, and gas stripping effectivity. These models are used as inputs for a bioprocess design and included in a final computational simulation of the chosen production process, as illustrated in figure 1, thereby highlighting the importance of tailoring biological parameters to process and product requirements.

**Figure 1:**
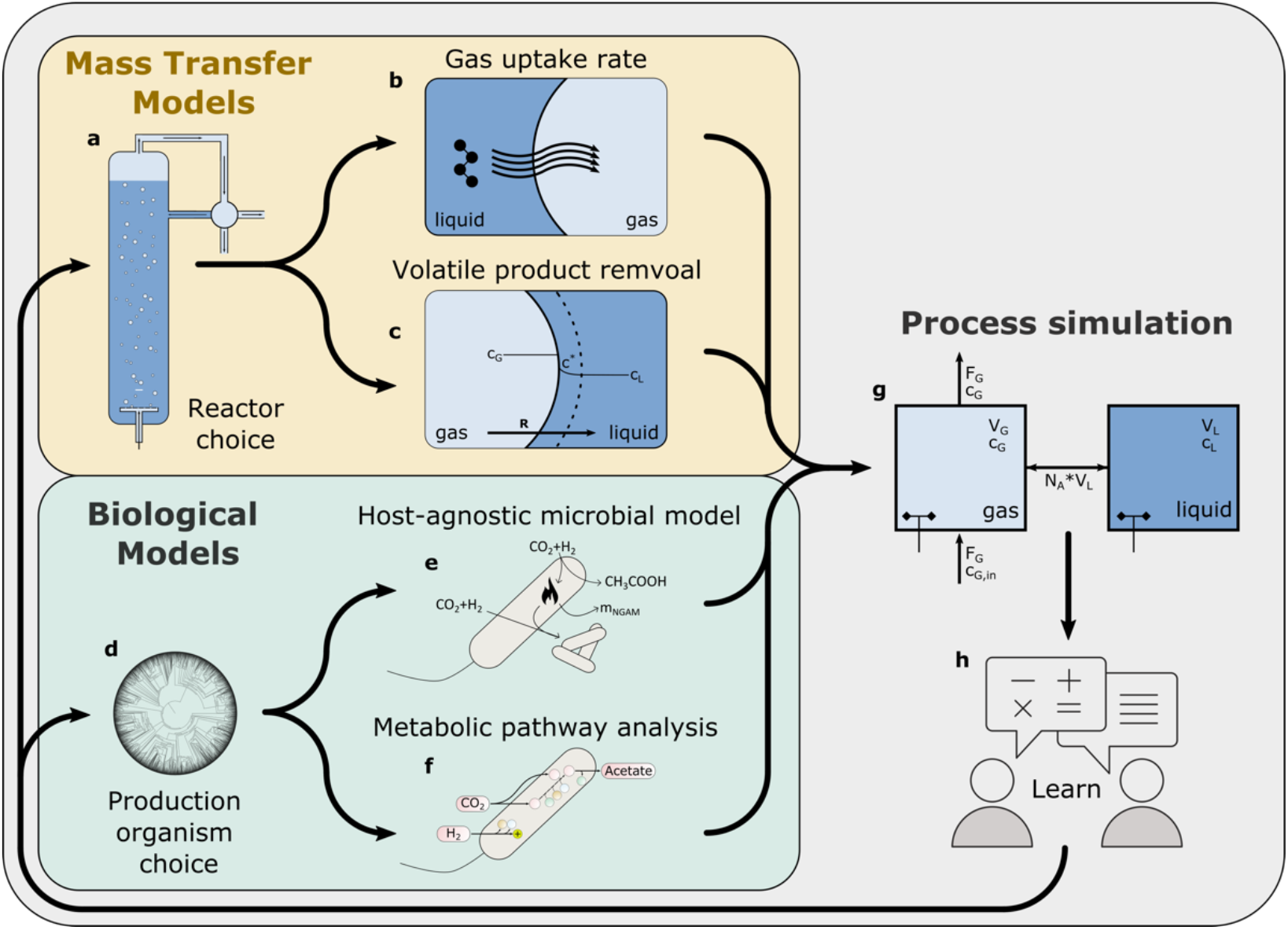
Overview of the models used in this work. Starting with a choice of reactor (a), mass transfer for both the gas substrate uptake rates (b) and the volatile product recovery are modeled (c). For a chosen production organism (d), a model based on general microbial growth, overall stoichiometry, and reaction thermodynamics is built (e), complemented with a more detailed stoichiometric pathway analysis (f). These models are used to run a process simulation (g) of which the learnings (h) are used to reevaluate the initial choices.

In bioprocess engineering, the choice of the host organism is often the first decision to be made. However, this choice is often limited to a few model organisms, or even more restrictive, to the specific organism that researchers have predominant experience with. While engineering might increase the performance of the specific organism under the desired conditions, better characteristics may be achievable by choosing a starting organism that is innately more suited to the envisioned product and hence required process. To capture this ambiguous starting point, and let process requirements dictate ensuing host selection, a host-agnostic bacterial model of microbial growth dynamics can be used, based only on universal overall reaction stoichiometry and general microbial growth properties^27^.

As summarized in Figure 1, we start with assessing the substrate solubilization by modeling the gas-liquid mass transfer rates through two-film mass transfer theory. We then investigate the effect of temperature on acetogenic growth using temperature dependent, host-agnostic black box biological models of acetogenic yields. These two models are then used in a process simulation for acetate production in an industrial-scale Bubble Column Reactor (BCR). To test the possibility of expanding the product portfolio beyond acetate, we model acetogenic metabolism in more detail using a stoichiometric model of the WLP, thereby comparing the energetic balance of different possible products. A comparison of vapor pressures for a range of VOCs tests the feasibility of *in-situ* product recovery.

## 2 Results

As a starting point of this study, initial process design considerations had to be made. For a bioprocess with gaseous substrate, mass transfer rates tend to be limiting. Reactor types with high gaseous mass transfers are thus suitable. BCRs are conceptually the simplest version of a high transfer rate reactor and are considered to be the benchmark for industrial scale processes reliant on efficient gas transfer^28^. They consist of a tall column with pressurized air being injected at the bottom and few moving parts. This simplicity allows for a bioprocess with low operation and maintenance costs. Other reactors, such as packed bed reactors^29^ or U-loops^30^ have even higher gas transfer rates^31^ but are more complex and costly and so were not included in the present work. Based on a thermodynamic and economic assessment of syngas fermentation by Redl et al.^32^ focused on maximizing gas uptake whilst remaining within practical limits, we decided to simulate a 30-meter-tall BCR with a radius of 3 meters, continuously fed with 10,000 m^3^/h of gas.

### 2.1 Gas-liquid mass transfer rates favor thermophilic conditions

While higher temperatures tend to favor mass transfer rates, they decrease gas solubility, making the interplay between the two difficult to predict. To investigate the overall effect of temperature on gas saturation concentrations and diffusion rates, the uptake rates of gaseous substrates into the fermentation broth were modeled. Two-film mass transfer theory^33^ was used to investigate the effects of temperature on the volumetric mass transfer rate (R) through the combined effects of saturation concentration (c*), volumetric mass transfer coefficient (k_L_a) and concentration of dissolved gas in the liquid. Both c* and k_L_a are temperature dependent and specific to the compound, but k_L_a is additionally also dependent on the reactor type and operation. Therefore, k_L_a is modeled for a large-scale BCR type as chosen and described above. Assuming full microbial conversion of all dissolved gasses, R and its components are expressed as functions of temperature and presented in Figure 2 (a-d).

**Figure 2:**
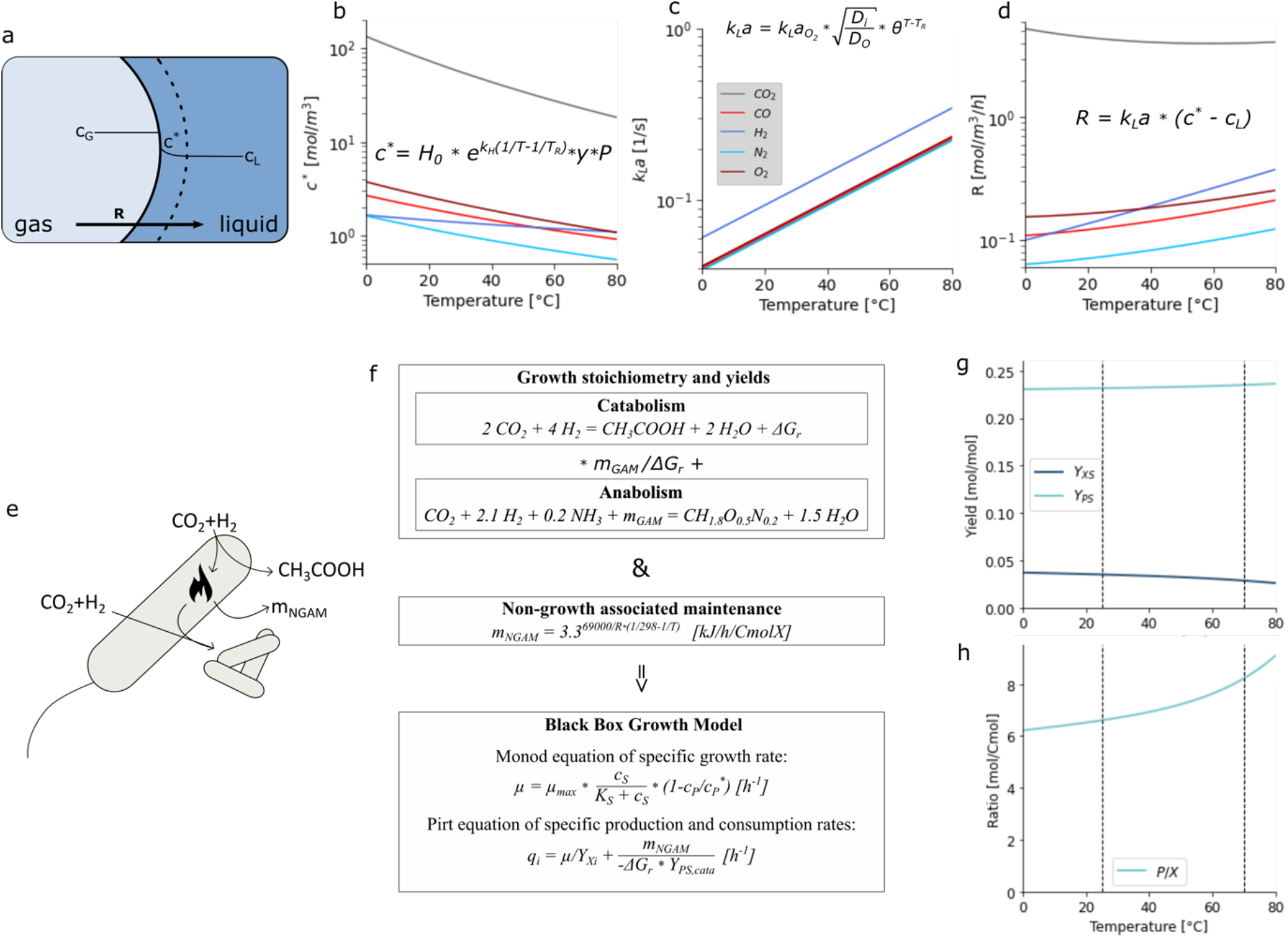
Mass transfer modeling of gaseous substrate uptake (a-d) and host agnostic black box microbial model (e-h) for CO_2_ fixation to acetate. Temperature-dependent changes in saturation concentrations (b), k_L_a (c), and the overall volumetric gas transfer rate (d) for CO_2_, CO, H_2_, N_2_ and O_2_, assuming gas transfer is limiting. Illustration of the black-box metabolic model (e), how it was built, including reaction thermodynamics, maintenance requirements, and maximum growth rate (f), and the resulting comparison of acetogenic growth and production yields on substrate (g) and product to biomass ratio (h) as a function of temperature.

While the saturation concentrations of all gasses decrease as temperature increases (Figure 2a), the volumetric mass transfer coefficient, increases exponentially (Figure 2b). Combining these two factors to calculate the overall gas transfer rate shows that the k_L_a impacts overall transfer rates most in the case of CO, H_2_, N_2_, and O_2_ (Figure 2c). The overall transfer rate of CO_2_ decreases, but only by 0.97-fold between 30 and 80 °C, remaining an order of magnitude higher than the other gasses. O_2_ and CO transfer rates increase by around 1.5-fold, while especially the transfer rate of H_2_ increases steeply, by 2.4-fold over the same temperature range. For H_2_, this is a combination of having the highest k_L_a and a saturation concentration least affected by temperature. Considering that H_2_ is the most expensive and hence often rate limiting gas in acetogenesis, this is an important improvement for overall CO_2_ fixation rates of the process. Thus, the volumetric mass transfer rate, and not the gas solubility, dictates the availability of the gaseous substrates, specifically favoring higher temperature operation.

With k_L_a being inherently dependent on the reactor type, size, and operation, these observations cannot be generalized to all setups. If the k_L_a values differ, their contribution to the volumetric gas transfer rate will change, which should be investigated per reactor type. However, with saturation concentration not being affected by a reactor choice, this effect would thus be even more pronounced for reactor types with higher mass transfer rates.

### 2.2 Temperature dependence of acetogenic yields

Thermodynamic effects are not limited to the physics of gas transfers, but also affect chemical reactions and accordingly, the metabolism of cell factories. Considering the metabolic diversity of acetogens, host agnostic modeling was here applied to capture population complexity, as often performed in microbial ecology or food preservation. To this end, the growth parameters of specific organisms are substituted by the maximum possible growth rate of microbes as a whole. It has been observed that both the growth rate and the death rate of such mixed samples follow the same exponential increase as the Arrhenius equation, used to model the temperature dependence of reaction rates^34–38^. The microbial death rate is conceptually a different way to account for the same biological phenomenon as the Non-Growth Associated Maintenance (NGAM), a parameter to account for yield variation with growth rates^27^. Thus, a temperature-dependent, host agnostic model was made, where the maximum growth rate (μ_max_), NGAM, and the growth and production yields that serve as inputs for Monod and Pirt kinetics^27^. The growth and production yields are deduced from the overall reaction stoichiometries, which in turn is predicted by balancing out the Gibbs free energy of reaction (ΔG_r_) generated by catabolism with the Growth-Associated Maintenance (GAM) (Figure 2f). As acetate is the main product from gas fermenting acetogens, we focus further efforts on this product, although this can be exchanged for other WLP products of interest.

With increasing temperature, the predicted product to biomass ratio increases by 14% between 30 °C and 60 °C (Figure 2h). Because ΔG_r_ decreases as a function of temperature, this observation is solely explained by the increased NGAM energy requirements at the higher temperature, as predicted by Thijhuis et al.^39^ (Supplementary Figure S1a, b). The product yields remain largely unchanged, with only a slight 1% increase between 30 and 60 °C, while the biomass yields decrease by 5-11% over the same range because of the increased maintenance requirement (Figure 2g). Therefore, increasing temperature steers substrate towards increased product formation, by favoring acetate over biomass production. It should be noted that this result is based on simplified metabolic energy demands that would require experimental confirmation. Studies on the effect of temperature on production to biomass yields of mixed acetogenic cultures, and on the metabolically closely related methanogens do confirm these predictions^40^.

The yield calculations show the maximum achievable theoretical yields based on thermodynamics, where in practice yields would be lower. Cells are limited in their energy generation capacity by intermediate molecules such as ATP or through ion gradients, thus lowering the achieved yields. Different acetogenic organisms differ in their energy and redox metabolisms^41^, regardless of their optimum growth temperature. Considering evolution and selection favors species that have efficient energy metabolism and hence growth at their given temperature optima, the experimental product and biomass yields would nevertheless follow the observed trend.

### 2.3 Process Simulation

To investigate the effect of the calculated yields and maintenance requirements in the BCR setup, whilst also accounting for the changing gas mass transfer rates, the black box model of acetogenic metabolism was included in a simulated BCR at both 30 and at 60 °C (Figure 3). In this simulation, the process is simplified by assuming a continuous product recovery, keeping acetate concentration in the liquid at zero. This assumption is necessary to investigate the steady state of a continuous production process, unhindered by product inhibition. *In situ* product removal can be achieved through classical separation techniques such as liquid-liquid extraction^42^, by using membranes^43^, or by converting the produced acetate into other, more easily separable products, such as VOCs in a coupled secondary fermentation^44^. A process presents many operation parameters that can be fine-tuned. For BCRs, volume, height-to-diameter ratio, flow rate, bubble size, and feed gas composition are key parameters. With all but the last already defined in the initial reactor setup, the main parameter left to define is the feed gas composition. If the gasses were fully dissolved, the ideal feed composition would be the same as the stoichiometric uptake ratio, namely 2:1 (H_2_:CO_2_). However, since the transfer rate of CO_2_ is an order of magnitude higher, H_2_ must be fed at greater amounts. To investigate this, we model the steady-state outcome as a function of the mol fraction of H_2_ in the feed gas (Figure 3a, b). The predicted optimal gas composition is then used as an input for following simulations.

**Figure 3:**
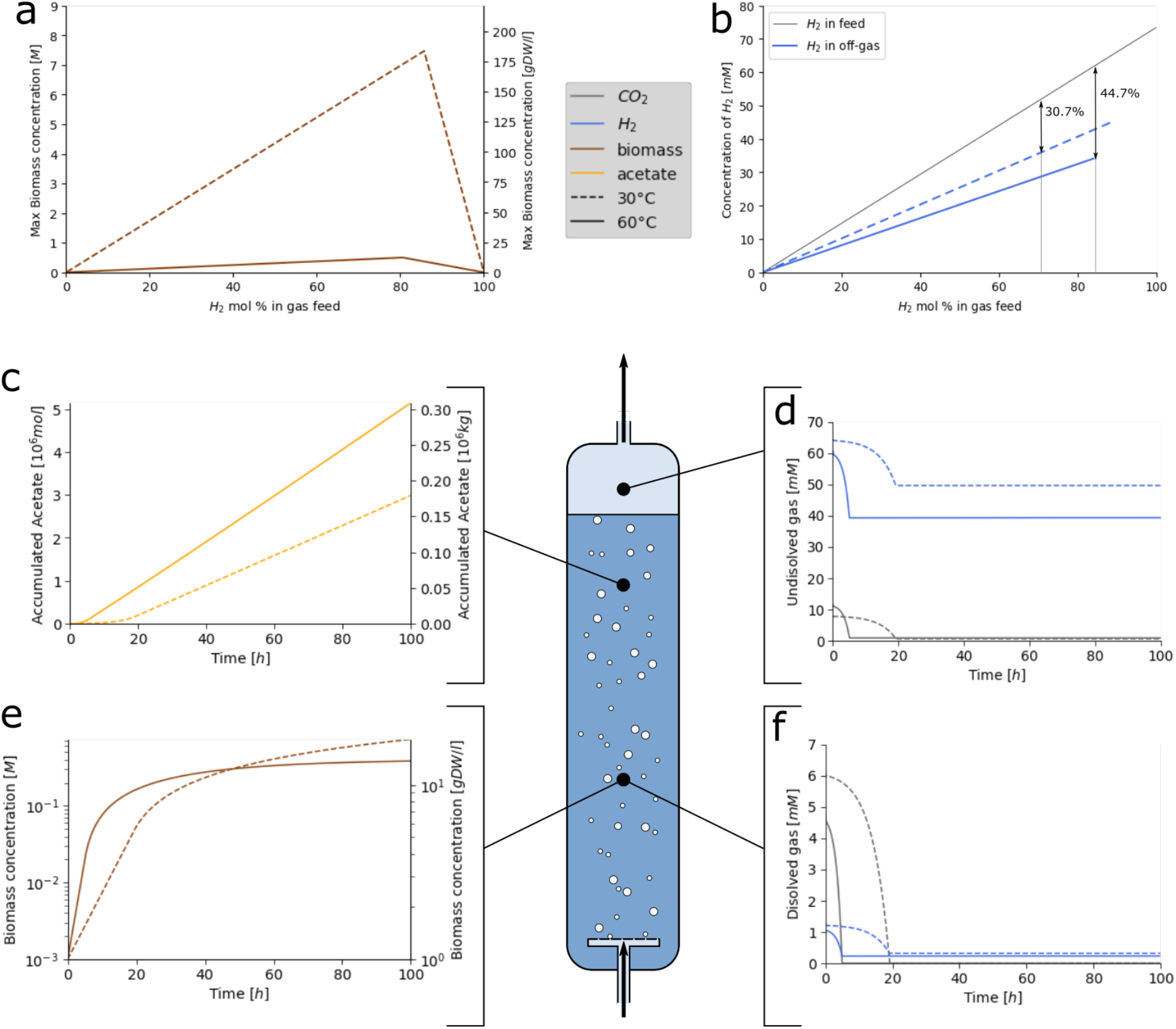
Steady-state analysis for optimal gas composition (a, b) to find the maximal reachable biomass concentration (a) and the ratio of H_2_ consumed from the feed (b) as functions of the mol fraction of H_2_ in the feed gas. The optimal gas composition is then used for a simulation of acetogenic growth in the described BCR at 30 °C and 60 °C (c, d, e & f) with continuous product removal. The results are split into accumulation of the product, acetate (c), undissolved gasses in the gas phase and off-gas (d), biomass concentration (e), and dissolved gasses in the fermentation broth (f). In all plots dashed lines show simulations at 30 °C and plain lines at 60 °C.

In the steady-state comparison (Figure 3a) the maximum reachable biomass initially increases with increasing H_2_ mol fraction, until it abruptly drops between 80 mol % and 90 mol %. At these points the H_2_ is no longer the limiting substrate, as CO_2_ becomes limiting. To compare how the increased H_2_ mol fraction in the feed gas is reflected in the off gas, another steady state analysis is done (Figure 3b). Because the percentage of H_2_ consumed from the feed gas, i.e. 30.7% at 30 °C and 44.7% at 60 °C, remains unchanged up to that point (Figure 3b), it is optimal to choose a gas composition as close as possible to this ratio, much closer to 9:1 (90 mol %) than to the stoichiometric 2:1 ratio (67 mol %). The exact values are found to be 0.86 mol % at 30 °C and 0.81 mol % at 60 °C, which are used for the respective gas compositions of the following simulations. It is noteworthy that, at 30 °C, the predicted maximum achievable biomass concentration reaches cell densities so high that they would increase fluid viscosity, which in turn decreases the k_L_a and so transfer rates. Overall, high cell density is to be avoided in BCRs for this reason, which is also an argument in favor of a thermophilic process^28^.

With the gas-fed, batch reactor simulations at 30 °C and 60 °C, temperature effects on yields, biological rates, and transfer rates can be calculated (Figure 3 c-f). Initially, dissolved gasses are at saturation concentrations, which are higher at lower temperatures, however, they begin to decrease as the cells grow (Figure 3f). A pseudo-steady state is reached where the mass transfer of H_2_ becomes limiting, keeping the concentration of dissolved gasses at 0 mM. During this pseudo-steady state, the concentration of undissolved gasses is lower at 60 °C, meaning that more gas is converted and less is wasted through the off-gas (Figure 3d). From a biological point of view, initial growth rates are at their theoretical maxima, which are higher at 60 °C. However, when gas transfer becomes limiting, the growth rate gradually decreases and the cells at the 30 °C condition would slowly grow to higher densities than those at 60 °C (Figure 3e). Maintenance energy requirements increase with temperature and energy is generated by acetate production; less biomass can thus in theory be sustained at 60 °C. The product formation rates on the other hand remain constant (Figure 3c), with 0.88 and 1.5 tons of acetate being produced in the reactor per hour at 30 °C and 60 °C respectively, corresponding to a specific productivity of 1.56 and 2.68 kg acetate per m^3^ per hour. Overall, the simulations predict 1.7-fold higher production rates at 60 °C. These amounts are in a comparable range to the estimate by Redl et al.^32^ of 3.79 tons of acetone produced per hour in the same reactor setup^32^. A decreasing biomass formation combined with a higher product formation pushes towards higher product on biomass yields at higher temperatures. These two simulations support the previous findings, thereby confirming that the transfer rate of a gas is more influential than its saturation concentration. Additionally, higher maintenance requirements at 60 °C result in an advantageously higher yield of product per biomass and production rates that are almost doubled.

### 2.4 Product comparison by stoichiometric modeling of the WLP

As the data thus far shows, temperature greatly affects the gas fermentation process, leading to advantageous properties of thermophilic production for catabolic products, specifically for CO_2_utilization. To develop a platform technology for CO_2_ consumption, modular production of various products of interest at high yield should be considered. Additionally, insightful perspectives on the metabolism of acetogenesis can be gained when comparing acetate production to various other possible products. Several studies show the heterologous expression of genes allowing for acetone and isopropanol production derived from acetyl-CoA by acetogens themselves^10,18,20,45–47^, expanding the list of small compounds that are innately produced by acetogens. With the constant development of synthetic biology tools, expanding this to thermophilic acetogens has become possible^20^, though difficult and time-consuming. Considering that the WLP is an energy demanding pathway relying on product formation for its energy, we investigated which common bulk biochemicals are energetically favorable metabolites to produce and thus suitable targets.

The simplest method to compare the thermodynamic feasibility of acetogenic carbon utilization into each of these various products is by calculating the difference in ΔG_r_. The reaction stoichiometry and ΔG_r_ were calculated for a range of possible products in the same way as for the black box model of acetogenesis (Figure 4b, Supplementary Table S2). Products considered are derivatives of acetate or acetyl-CoA, namely: ethanol, butanol, acetone, butyrate, and butanediol. Interestingly, all selected products have similar ΔG_r_. To determine for each product how much of that ΔG_r_ is usable for the cell’s metabolism, a stoichiometric pathway model, focusing on the energy and redox intermediates was constructed (Supplementary Figure S3). The metabolic map can be divided around its core metabolite acetyl-CoA, yielding the upstream WLP and downstream product formation (Figure 4a). To deduce the overall stoichiometry for each of the substrate and product combinations, the carbon incorporation to acetyl-CoA and the following product formation can be summed up. Accordingly, acetate and ethanol are the only products that fully restore the ATP consumed upstream to produce acetyl-CoA through their downstream metabolism, directly from substrate level phosphorylation (Figure 4a, stoichiometric sums in Supplementary table S1). Acetone and butyrate restore half, whereas butanol and butanediol do not generate any ATP. While ATP is the main biological energy carrier, energy can also be stored in redox intermediates such as NADH, NADPH, or ferredoxin, or as sodium or proton gradients across the cell membrane, which can be converted to ATP through a sodium- or proton-driven ATP synthase. Although central to the cell’s metabolisms, these differ fundamentally between acetogenic species^41^. For some acetogenic species, these mechanisms have been well defined. *C. ljungdahlii*, which was used as a template for the stoichiometric model, uses three of these enzyme complexes. To form ATP from a proton gradient over the cellular membrane, it uses a proton-driven ATP synthase. Additionally, ferredoxin NADH-linked hydrogenase splits H_2_ to reduce ferredoxin and NAD^+^. Lastly, Ferredoxin:NAD^+^ Oxidoreductase (Rnf) allows NAD/NADH to reduce ferredoxin and vice versa, also using the proton gradient as a driving force. When including these reactions in the stoichiometric model, the proton and redox factor requirements or production of the metabolic reactions can be translated to ATP generation only, which determines whether the pathways can generate energy for cellular growth (Figure 4b, Supplementary Table S2).

**Figure 4:**
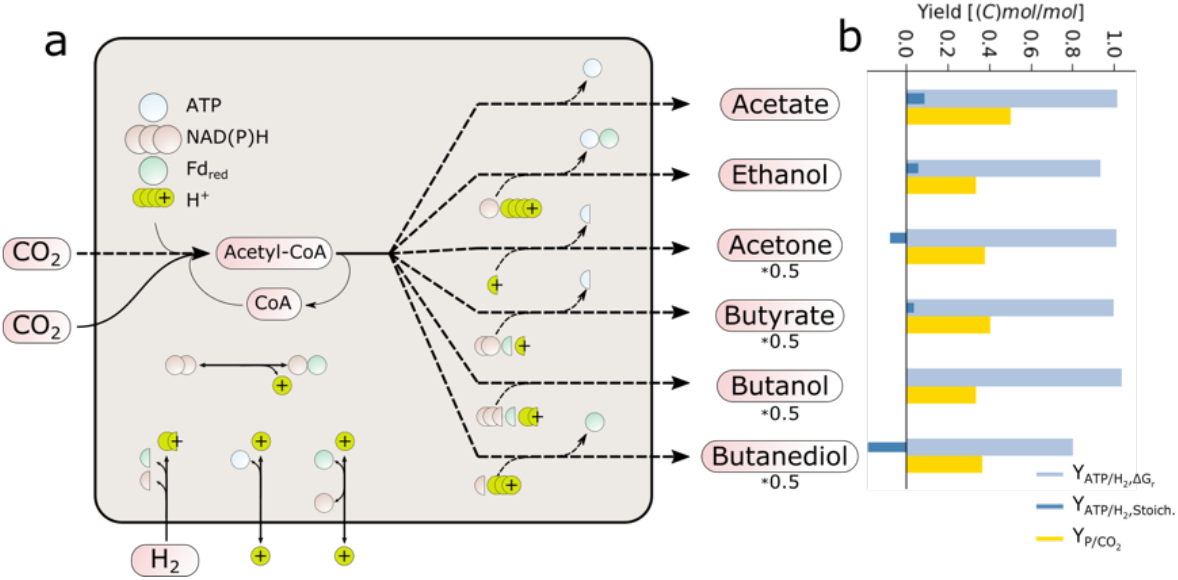
Product comparison by metabolic pathway analysis of the WLP showing a simplified pathway map of the WLP and its various products (a), and the ATP and product yields for different acetogenic products (b), as predicted by reaction thermodynamics (light blue) and how much of that theoretical maximum is stored in ATP as predicted by the stoichiometric model (dark blue). The product per substrate yields (yellow) is a metric for how much carbon from the CO_2_ is fixated into the products. ATP yields are shown in mol/mol, whereas product yields are shown as Cmol/Cmol.

When comparing potential fermentation products, two metrics are especially important. Firstly, the ATP yield on the electron donor determines how much energy the cells can generate for both growth and maintenance. Secondly, the product yield quantifies how much product is formed per CO_2_. In general, a rough trend of decreasing yields with increasing product size can be observed, with acids having higher yields than alcohols. Acetate stands out through both its ATP and product yield. Compared to ethanol, which is a downstream product of acetate, it is more beneficial for two reasons. Firstly, the reaction of acetate to ethanol through acetaldehyde requires two reducing equivalents. Secondly, the acetate proton symporter feeds the proton motive force in a wide range of organisms, including acetogens^48^. The proton motive force can power ATP synthase and, in some acetogens including *C. ljungdahlii*, the bifurcating enzymes such as Rnf. Butanediol gives the lowest ATP yield, requiring ATP to utilize CO_2_ as a substrate. Acetone is also energetically unfavorable, requiring ATP to be produced from CO_2_. An alternative acetone production pathway has recently been proposed, which directly generates ATP by using a phosphate butyryltransferase and a butyrate kinase enzyme that will promiscuously catalyze the transformation of acetoacetyl-CoA to acetoacetate, through acetoacetyl-P^49^. However, when including this pathway in the stoichiometric model and comparing the yields to the pathway from *C. acetobutylicum*, the same overall yields are observed, meaning that the pathways are energetically equivalent (Supplementary Figure S4). Negative or neutral ATP yield values do not mean that the products cannot be formed, but that flux towards another product that does generate energy is required to supply the required energy for both production and growth. This is also a common phenomenon when strains are heavily engineered past energetic limits to force formation of a specific product, resulting in significant byproduct formation to meet cellular energy and redox demands.

The stoichiometric model builds on a steady state assumption, so all accumulations except substrate, product, and ADP/ATP are set to be zero. *In vivo*, the redox intermediates NADH, NADPH, and ferredoxin are involved in other parts of cellular metabolism. Modeling a steady state pushes flux through the bifurcating enzymes, which could affect ATP yields. Therefore, one should be cautious when determining a clear cut-off value for the feasibility of producing compounds. Nonetheless, the combined properties of acetate, having both a high ATP and product yield make it a suitable candidate for acetogenic carbon utilization. Evolutionary pressure will push for its stable production, and its production results in the most carbon being utilized.

### 2.5 Volatile product comparison for *in situ* recovery

Having demonstrated that thermophilic fermentations are theoretically beneficial for the uptake and conversion of gaseous substrates, one can also consider another advantage of increased temperature, specifically the possibility of *in situ* removal of volatile products of interest through the off-gas. This enables more simplified downstream processing, prevents product toxicity to production strains and feedback inhibition on production pathways, overall leading to higher specific productivities of a process. Low boiling point derivatives of acetate, such as acetone, previously produced as a product of the WLP^20^, are possible products to consider. Evaporation can happen either by gas stripping, which depends on its vapor pressure, or by boiling, which happens when the vapor pressure exceeds the ambient pressure. Thus, the vapor pressures of selected volatile compounds were expressed as a function of temperature for a range of possible products (Figure 5a). Of all modeled compounds, acetone is the most volatile, with a boiling point below 60 °C at atmospheric pressure (Figure 5a). Other products such as ethanol and butanol closely follow, while most others are around or below the vapor pressure of water. However, the pressure in the reactor is not atmospheric as it varies significantly by the depth in the broth, especially for tall BCRs (Figure 5a). From here, one can observe the depth at which the pure compound of interest will boil at the given temperature. Assuming a process temperature falling within the physiologically relevant 30 °C and 70 °C range, evaporation by boiling would only occur for pure acetone in a fraction of the reactor around or above 60 °C.

**Figure 5:**
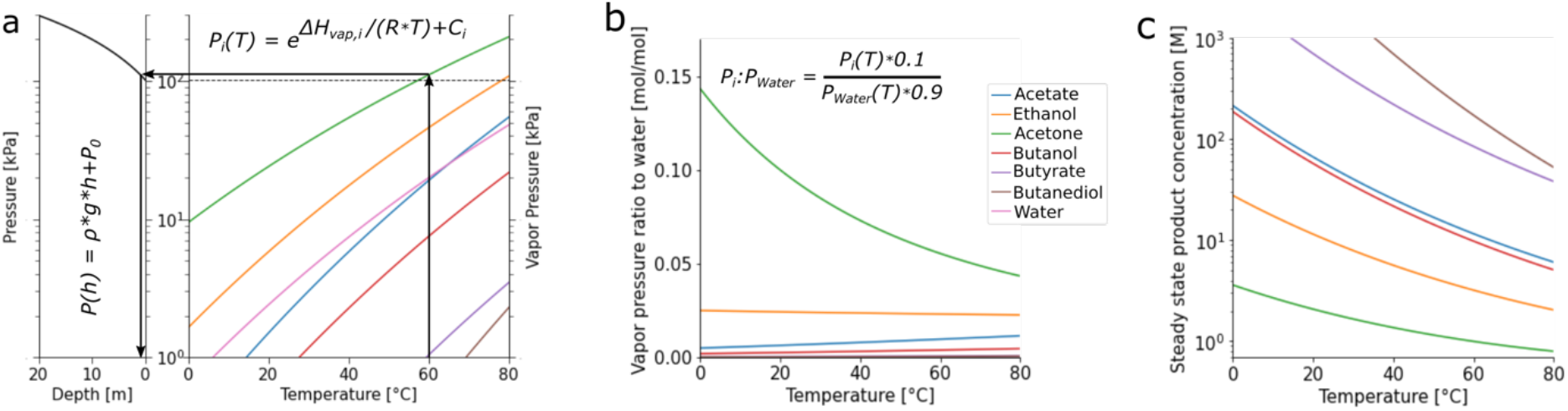
Analysis of volatile product recovery, represented through investigation of the vapor pressures of selected pure solutions of VOCs as a function of temperature (a, right), juxtaposed to the increase of pressure as a function of depth in the reactor (a, left). The arrows show how both can jointly be read to find the reactor depth at which a chosen compound reaches its boiling point for any reaction temperature. The ratio of vapor pressure of VOC to water in the off gas, for biologically relevant concentrations of 1% mol fraction VOC in aqueous broth determined across a temperature range (b). The product concentration in the bulk of the BCR at steady state as a function of temperature, based on the mass transfer models and overall reaction stoichiometry (c).

Additionally, in a biological fermentation process where cells are suspended in liquid media, one would expect an aqueous broth with product concentrations at a maximum of 5-10% (w/v), in part due to product inhibition and toxicity^50–52^. In an aqueous solution, both the boiling point of the solution and the vapor pressure of the solute change. The change in the boiling point of a mixture depends on complex physicochemical interaction properties of the solvent and solute, which must be experimentally determined. For the example of acetone, studies have experimentally measured these values for aqueous mixtures at varying mol fractions and pressures and identified that the boiling point of the mixtures starts at that of pure acetone, remains relatively flat until a 10% acetone mol fraction, and then steeply increases to the boiling point of pure water^53^. In a BCR setting, even at thermophilic temperatures, boiling is thus likely insufficient to allow effective product recovery. Yet, even below the boiling point, the VOC can still diffuse to the gas phase through the exerted vapor pressure. Gas stripping could therefore provide a second evaporative mechanism for *in situ* product recovery. As with boiling, this needs to be corrected for the mole fraction of the product in the solution. In solution, the partial vapor pressure can linearly be related to the molar fraction of the compound and the vapor pressure of the pure component through Raoult’s law. For acetone, the most volatile and hence favorable compound, assuming a biologically relevant mol fraction of 1% (corresponding to a relatively high concentration of approximately 0.6 mol/L or 32 g/L), the vapor pressure would be 1% of that of the pure solution, pushing it far below that of water. Hence, at a given temperature, the ratio of VOC to water that will evaporate, at steady state, equals the vapor pressure of the solute divided by that of the whole solution (Figure 5b). With higher temperature, the ratios change as the vapor pressures of the different compounds change independently. The vapor pressure of water increases more steeply with temperature than most dilute VOCs (Figure 5a). In the example of a 1% solution of acetone, the ratio decreases from 7.6 to 5.5% (Figure 5b) between 37 °C and 60 °C, but still corresponds to a 7.6 and 5.5-fold enrichment of the VOC from the liquid to the gas.

The feasibility of volatile product recovery relies on the ability of the gas stripping to push down the product concentration to avoid inhibitory effects and allow continuous production. We thus chose to investigate for the given BCR at the given feed rate, whether the exerted vapor pressure is sufficient to keep the product concentration low (Figure 5c). At a steady state with continuous production, there is no product accumulation, so product removal rate must equal the production rate. Using both the mass transfer rate of the substrate, H_2_, and the overall reaction stoichiometry for each product, the maximum product formation rates are calculated. Assuming that the product concentration in the off-gas reaches its saturation, the concentration in the bulk is in equilibrium with the concentration in the gas, as dictated by the vapor pressure. The two products with the lowest predicted concentrations are ethanol and acetone. At 60 °C, ethanol is predicted to reach a concentration of 3.2 M (147.2 g/l), while acetone reaches 1.0 M (57.8 g/l). Only acetone comes to a biologically relevant 5-10% w/v product concentration range. However, tolerance is highly host specific, and only once a host is selected is it possible to confirm the tolerance levels. If needed, tolerance can be improved with strain engineering or laboratory evolution^54^. As opposed to the two-film theory model used for the absorption model, the product removal calculations are built only on steady-state concentrations dictated solely by the compound’s vapor pressures. It is not tested whether a steady state is reached and the gas is saturated before it leaves the reactor. This implies that the molar percentage of product in the off-gas could be higher if the transfer rate of the product is higher than that of water.

In conclusion, while operating above the boiling point of the pure product does not guarantee successful *in situ* product removal, gas stripping could be sufficient to continuously recover acetone in a BCR setup at thermophilic conditions and keep a concentration within a biologically relevant range. There is no clear temperature threshold above which the recovery becomes feasible, rather, the higher the temperature, the lower the steady state product concentration is and the less product inhibition affects the process. Inevitably, continuous product evaporation will also evaporate large amounts of water, which forms a challenge for the process design and operation.

## 3 Discussion

Microbial gas fermentation is an efficient platform that enables the conversion of CO_2_-rich waste gas streams into biomass and organic carbon compounds, which can be used as sustainable products. To harness this power, optimization of both the production strain and process are critical to achieve sufficiently high efficiencies and productivities, to outcompete petrochemical production in the economic-incentive-driven society.

Applying host-agnostic mechanistic models allows for tailored selection of microbial hosts based on final process requirements. Particularly advantageous to using host-agnostic black box modeling is the low dependency on large *a priori* experimental datasets, which are often not available when pushing the boundaries of novel production processes and microbial phenotypes. In this way, a more efficient allocation of resources can be achieved when process and metabolic modeling are used to narrow the solution space that must be experimentally probed. Requiring limited data input, this approach offers a modular platform to explore novel bio-manufacturing possibilities, moving the field away from traditional production organisms and processes. Although such modelling approaches are advantageous to limit the experimental test space, they can fall short of capturing intricate processes, either because they are a simplification of complex systems, or because certain factors are unknown. Therefore, critical biological and experimental aspects presented in this work should be considered and validated. Nonetheless, this approach could be applied to query other unconventional types of metabolism, such as the many other types of anaerobic respiration^55^.

We show that for catabolic products, product yields increase with temperature due to the higher demand for maintenance energy. Specifically, for acetogenesis, metabolic modeling performed here shows that the energetic feasibility of acetogenesis depends solely on the downstream metabolism of product formation, to compensate the energy-demanding WLP, with acetate being the most energetically favorable product whilst also utilizing the most carbon per electron donor. When other products are desired, lower yields would be achieved, as by-product formation is required for energy generation. This is often what is observed in gas fermenting organisms^15,56^, resulting in more complex downstream processing and overall lower titers.

Using mass transfer models and process simulations, we have investigated improved operating conditions for a BCR. We demonstrate in this study that higher temperatures are beneficial for gas uptake rates, essential for substrate availability in acetogenesis, while to date predominant industrial work with gas fermentation is performed in the mesophilic temperature range^57^. While BCRs are a good starting point for gas fermentations due to their simplicity, scalability, and low maintenance^58^, other reactor types can also be envisioned. Extensive modeling and experimental studies of, for example, continuous stirred tank^59^, packed bed^29^, or external loop^30^ reactors have also been demonstrated. With solely gaseous substrates, gas transfer is expected to be limiting regardless of the reactor type. Considering that the main difference between the reactor setups from a mass transfer perspective is the value of k_L_a^60^, the findings of this research can also be applied to other reactor types. To our knowledge, there is currently no quantitative comparison of reactor performance between setups for the specific purpose of gas fermentation. This could provide valuable insights and further improvements into gas fermentation-mediated CO_2_ upcycling.

Gas transfer rates are only limiting as long as the fermentation is substrate limiting. If, through product accumulation, growth becomes product inhibited, product removal becomes the bottleneck. Then, *in situ* product removal can greatly benefit specific productivities. However, our calculations predict that product removal is only within reach for acetone and comes with the challenge of predicted dilute concentrations in the recovered gas, due to the large amounts of water evaporating. Other studies have also highlighted this challenge for other types of fermentation processes such as ABE fermentation^61–67^. When comparing the thermophilic gas fermentation to these mainly mesophilic ABE fermentations, the former comes with the advantage of gas feed already serving as the stripping gas, while the ABE fermentation uses a dissolved substrate, meaning that a separate stripping gas needs to be used, adding to the process complexity and operation cost. Additional processing steps can help the purification of the volatile product. The simplest addition is a second distillation after condensation of the off-gas, possibly using high osmolarity to salt out the product^65^. This is however purely downstream and does not affect the product concentration in the broth. If experimental work would show that the concentration needs to be pushed even lower, an option would be to increase the gas flow by adding a stripping gas. Increasing the overall gas flow in a BCR can break the flow of the air bubbles and cause a relative decrease of k_L_a^28^, therefore the feed rate can be kept the same while changing the gas composition to include N_2_ or more CO_2_. More elaborate *in situ* product removal methods that do not rely on evaporation can also be considered, such as the use of membranes^43^ or polymeric resins^68^ that separate the product based on other physiochemical properties, such as polarity. These are much more complex and come with a whole new set of challenges, especially when used in the relatively dirty environment of a fermentation.

Overall, a BCR enables high gas absorption rates with minimal operation complexity and costs. Increased operation temperature favor both gaseous substrate uptake and formation rates and yields of catabolic products. Energy generation in acetogenic gas fermentation relies on product formation, with acetate being by far the favored product. However, acetate is a product of limited value that cannot be recovered *in situ* by gas stripping. To reconcile this with the found benefits of a thermophilic gas fermentation in a BCR, several strategies can be considered. A co-substrate could be added to expand the product range. Other products could be formed by acetogenesis despite their yields being less suitable, most likely producing a mixture of compounds. Acetate could be used as an intermediate metabolite for further conversion into more valuable compounds, either by chemical catalysis or by a second fermentation step by a heterotrophic production host.

Taken together, this work highlights how the interplay of process and metabolic modeling can be used to steer bioprocess design to harness novel phenotypes and to allow probing of process parameters such as temperature and feed composition. Starting from the final production aim, we work our way back to choose suitable reactor operation conditions and a fitting production host. Specifically, the temperature-dependent, host-agnostic growth model is a powerful novel tool to assess and compare possible production hosts when the reactor conditions are not yet known. Combined with mass transfer models and metabolic pathway analysis into a process simulation, this approach has allowed to quantitively compare fermentation conditions and narrow down the range of process possibilities. Here, the study is specifically applied to microbial CO_2_ fixation, aiming to assess the technology and push it further with the aim of providing a new tool to battle one of the most pressing global issues. Overall, this pushes the paradigm to rely less on conventional production organisms and organic carbon substrates, highlighting the possibility of efficiently converting gaseous waste streams into valuable chemicals, needed to achieve the ambitious climate goals society has set.

## 4 Materials & Methods

In the presented work, we combine thermodynamics-based reactor modeling, building upon the findings of Redl, et al.^32^, with steady-state stoichiometric modeling of metabolism. All calculations and simulations were done in Jupyter Notebooks, freely available on GitHub (https://github.com/biosustain/CO2_fixation_models/releases/tag/v3.0), using python 3.8, extended with the Numpy^69^, SciPy^70^, and Pandas^71^ packages, and Matplotlib^72^ for plotting. Additional packages used are listed in specific sections below. Illustrations are made with the open-source vector image software Inkscape.

### 4.1 Reactor properties

The reactor setup used in the simulations is a 30-meter-tall BCR with a radius of 3 meters, filled to 2/3 of its volume^32^. It is continuously fed with a mix of H_2_ and CO_2_. The feed rate is adjusted to obtain a superficial gas velocity (v^c^_gs_)^73^ of 0.1 m/s, in the range of conventional values for large scale BCRs^32,74^. The reactor is modeled as two ideally mixed entities, the gas phase, which represents all gas bubbles passing through the column, and the broth. Only the gas phase has a continuous flow-through. Exchanges between both phases are modeled as a single transfer rate for each compound. Unless specified otherwise, the temperature range used in the various aspects modeled is between 0-80 °C, corresponding to 273.15-353.15K. Further parameters, including the liquid volume (V_L_), the gas holdup fraction (ε)^28^, the pressure at the bottom of the tank (P_b_), and the logarithmic mean pressure (P_m_)^73^ were defined from the basic properties of the reactor (eq. 2-6). Constants and physio-chemical values are named in the nomenclature and their values are listed with their sources in Supplementary Tables S3, S4, and S5.

Superficial gas velocity:

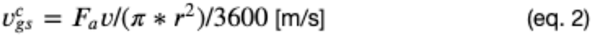

Liquid volume, assuming 2/3 filled:

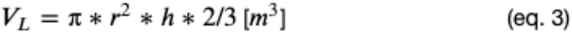

Gas holdup fraction:

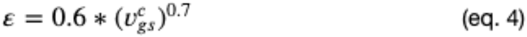

Pressure at the bottom, assuming 2/3 filled:

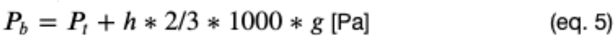

Logarithmic mean pressure:

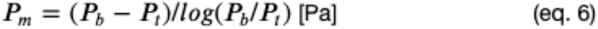

### 4.2 Gas-liquid mass transfer

To address the inflow and outflow terms of the mass balance, the mass transfer rates were determined for each compound across the gas-liquid phase boundary using the two-film theory^33,60^. In two-film theory, the division between two phases is seen as a boundary where each phase forms a film providing resistance to the mass transfer. The mass transfer through each of the boundary films (R) was expressed for each compound as a function of a coefficient (k), the surface area of the interface (a), and the concentration difference between the bulk of the phase and the interface. For gases that are poorly soluble in water, the liquid-side film resistance was assumed to be dominant, and the gas-side resistance negligible. This was done for CO_2_, CO, H_2_, O_2_, and N_2_ (notebook 01). The factors k and a were combined into a single term: k_L_a. The transfer rate is therefore a function of the volumetric mass transfer coefficient k_L_a, the saturation concentration, and the dissolved concentration in the liquid (eq. 7). The value of k_L_a is specific to both the reactor and the compound. For a coarse bubble system in the dimension range of the envisioned BCR, k_L_a of O_2_ at 20 °C was expressed as a function of the superficial gas velocity^28^. The k_L_a of all other compounds was extrapolated from that O_2_ using the diffusion coefficient of the new compound and the reference^75^ (eq. 8). Furthermore, k_L_a was expressed as a function of temperature by multiplying by the correction factor θ, to the power of the temperature difference (eq. 9)^28^.

The concentrations of dissolved gases at the liquid side of the interface were assumed to be at equilibrium, thus equal to the saturation concentration of the gas in water. This saturation concentration (c*) was expressed according to Henry’s law, as a function of a compound- and temperature-dependent constant (H_T_), the mol fraction in the gas, and the pressure (p), for which the mean logarithmic pressure is used^76^ (eq. 10). H_T_ was expressed as temperature-dependent expression, containing a known value of H at a given reference temperature, a compound-specific temperature correction factor k_h_ based on the van ‘t Hoff equation, and the reference temperature^76^ (eq. 11).

Transfer rate:

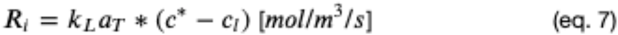

Volumetric mass transfer coefficient:

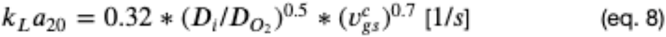

Temperature corrected volumetric mass transfer coefficient:

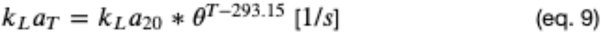

Saturation concentration

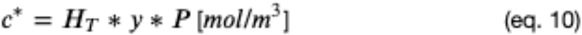

Temperature-corrected Henry’s law constant:

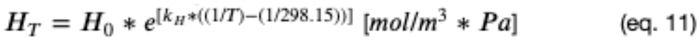

### 4.3 Acetogen thermodynamic black box model

Acetogenic yields were deduced from the overall reaction stoichiometries. These are established by determining the substrates and products and balancing the obtained equations^77^. Two reactions were defined, the catabolic reaction producing acetic acid (eq. 19) and the anabolic reaction producing biomass (eq. 21). The biomass composition was assumed to be CH_1.8_O_0.5_N_0.278_. To meet the nitrogen requirements of the anabolic reactions, ammonia (NH_3_) was included as a substrate.

Catabolic reaction stoichiometries:

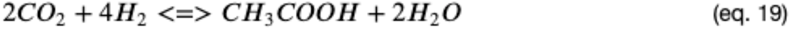

Anabolic reaction stoichimetries:

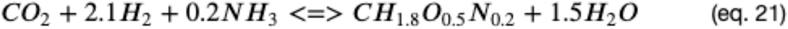

With the anabolic and catabolic reactions determined, the energy generated by catabolism and energy needed for anabolism were determined. Firstly, to determine the energy generated by catabolism, the Gibbs free energy of reaction (ΔG_r_) was calculated as a temperature-dependent function based on the Gibbs–Helmholtz equation (eq. 25). This equation corrects the Gibbs free energy of reaction by temperature, based on the Gibbs free energies of reaction and reaction enthalpy at standard conditions 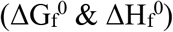, both being the difference of the Gibbs free energies and enthalpies of the products and substrates (eq. 23-24). Secondly, the anabolic energy requirement corresponding to the GAM was calculated. For autotrophic growth, GAM depends primarily on the nature of the electron donor^78^. It was thus estimated by extrapolation from other organisms to be 1000 kJ/Cmol_biomass_^78^. The overall reaction stoichiometries were predicted for any given temperature by finding the factor by which to multiply the catabolic reaction so that it generates the 1000 kJ that is required for the growth of 1 Cmol_biomass_ (eq. 26). These stoichiometries were then used to determine the yields of product on substrate (Y_PS_), biomass on substrate (Y_XS_), etc. by dividing the respective stoichiometric factors as ratios. NGAM is not included in the yields, as it is dependent on the specific growth rate. It is accounted for later, in the black box model.

Gibbs free energy of reaction:

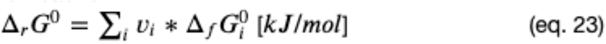

Reaction enthalpy:

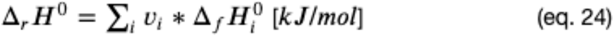

Gibbs-Helmoltz equation:

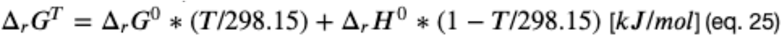

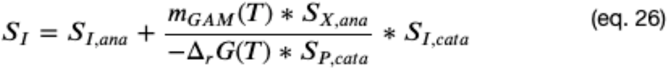

Building on the calculations above, a black box model was created for hydrogenotrophic growth. The model used here consists only of growth, consumption, and production rates, assuming a single rate-limiting substrate, namely the electron donor, H_2_. The specific growth rate is modeled using the Monod equation^79^, with a product inhibition term (eq. 28)^80^. Values for the saturation constant (K_S_) and the inhibitory product concentration (c_P_*) were obtained from literature or estimated (Supplementary Table S7). The maximum growth rate of acetogens (μ_max_) was predicted as a function of temperature with the Arrhenius equation (eq. 27), a model inspired by reaction kinetics, used in ecology and food safety^34,35,37,38,81^. The equation is a function of temperature containing three parameters: the gas constant (R), an equation-specific constant (A), and a substitute for the activation energy (B). To determine these parameter values, the function was fit to the maximum growth rates of 34 known acetogens found in literature, plotted by their optimum growth temperature. These values are listed in Supplementary Table S8 and were fitted using SciPy (Supplementary Figure S1C), which set the parameters to be A = 8.66 and B = 24166.93.

The Arrhenius equation for max growth rate:

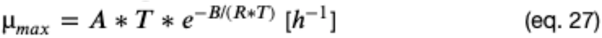

Specific substrate consumption rates follow Pirt kinetics and the same applies for product formation, as long as the product is formed in the catabolic reaction (eq. 29)^27^, with the maintenance term (m_i_) defining how much product or substate is formed or consumed through additional catabolic reaction needed to satisfy NGAM energy requirements (eq. 30).

Monod Equation, specific growth rate:

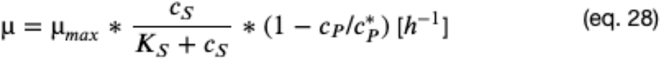

Pirt kinetics, specific consumption and production rate:

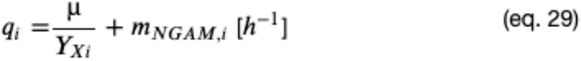

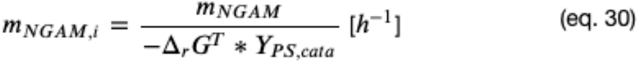

Similar to the maximum growth rate, NGAM correlates with temperature. Based on a wide range of studied anaerobic organisms, NGAM can be estimated through the temperature-dependent expression based on the Arrhenius expression, as proposed by Tijhuis et al.^39^ (eq. 31). This equation is compared to a new fit to their data in Supplementary Figure S1B.

NGAM:

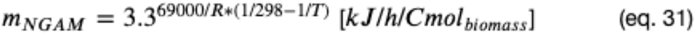

To assess and compare the energy generated to produce a wider variety of compounds, the catabolic reaction was determined using the same method as for acetate. With the reaction stoichiometry, ΔG_r_ is the difference of ΔG_f_ of the products and the substrates (notebook S02) for acetate, acetone, ethanol, propanol, butanol, and butyrate^77^.

### 4.4 Reactor Simulation

To simulate the change for each of the modeled compounds over time, a mass balance is set up over the system boundaries, either being the gas phase, liquid phase, or both (box 1). This mass balance defines the change in concentration as the sum of inflow, outflow, production, and consumption rates^60^.

#### Box 1

Mass Balances; Accumulation = In - Out + production - Consumption

Biomass: 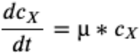

Acetate: 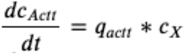

CO_2_, dissolved: 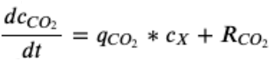

CO, dissolved: 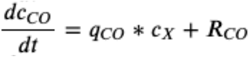

H_2_, dissolved: 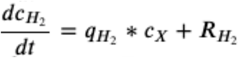

CO_2_, gas: 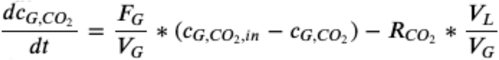

CO, gas: 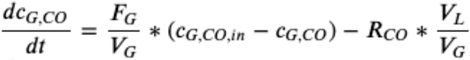

H_2_, gas: 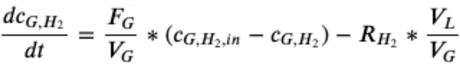

To find the optimal operation setting, the mass balances are solved at steady state, where the accumulation is null. Starting with the mass balance over the substrates in the gas phase and assuming that mass transfer is limiting, thus that its dissolved concentration is null, the gas concentration at steady state can be expressed as functions of the feed concentration (Box 2, notebook 03). Similarly, one can predict the maximal achievable biomass concentration, which could be reached at steady state. At this biomass concentration, all available substrate is required to satisfy maintenance requirements.

#### Box 2

derivation of steady state equilibria

##### 2.1 Optimal gas feed composition

An equation for *c*_*G, i*_ at steady state can be deduced starting with the gas phase mass balance.

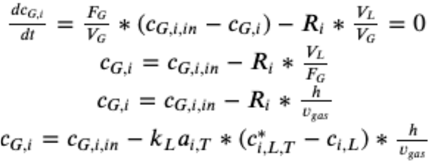

For the limiting substrate *i*, at steady state, where all of it is consumed in the liquid (i.e. *c*_*i,L*=_ 0), we have:

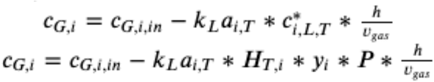

Using the ideal gas law, Raoult’s law and yields:

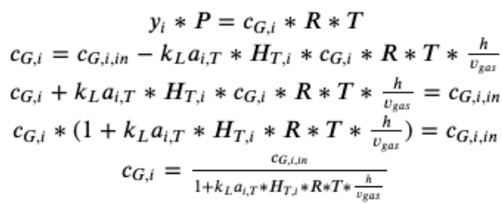

##### 2.2 Maximum steady-state biomass concentration

Using the liquid phase mass balance, we predict the maximum achievable biomass concentration, Where all subscribe is required for maintenance.

Mass balance of compound i in the liquid, with μ = 0:

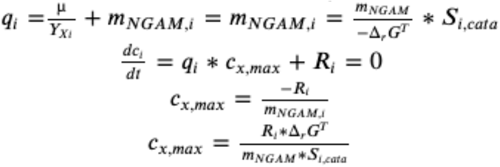

Plotting these results makes it possible to find the tipping point where the limiting substrate changes, namely the intersection of the defined functions for CO_2_ and H_2_. These points change with temperature as described in the results section. The gas feed composition at these tipping points was used as a parameter for further simulations. To simulate the reactor run at 30 °C and 60 °C, the set of differential equations (Box 1) was integrated using SciPy, with given initial conditions, and defined functions for growth (μ), production and consumption rates (q), and mass transfer rates (R) (notebook 04).

### 4.5 Volatile product recovery

To model the transfer of compounds from the liquid to the gas phase, namely evaporation of possible volatile products, the boiling points, and diffusion rates were expressed for each compound as functions of temperature. The vapor pressure is key for calculating both, it determines the saturation concentration in the gas for modeling diffusion using two-film theory and sets the boiling point as the temperature where the vapor pressure exceeds ambient pressure. In this analysis, acetate, methanol, ethanol, propanol, butanol, butanone, acetone, butyrate, and butanediol were included and water was used as a reference. Vapor pressure is temperature dependent and was thus defined for any compound as a function of temperature, using either the Antoine equation^82^ (eq. 12 & 13) or the Clausius-Clapeyron equation^83^ (eq. 14), or an approximation thereof^82^ (eq. 15 & 16). The gas was assumed to be ideally mixed and fully saturated by the time it leaves the column.

Antoine equation:

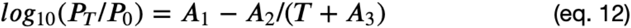

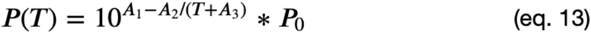

Clausius-Clapeyron equation:

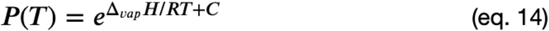

Clausius-Clapeyron approximation:

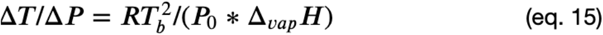

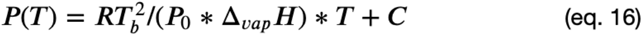

When compared, the Antoine and Clausius-Clapeyron equations yield highly similar curves, while the approximation is only good for temperatures within 10-15 °C of boiling point (notebook S01, Supplementary Figure S2). Therefore, the Clausius-Clapeyron equation was used, which is preferred over the Antoine equation because it requires fewer parameter inputs. Indeed, the pressure can be temperature corrected based only on the molar enthalpy of vaporization (ΔH_vap_) and a constant C, which is first fitted from a known vapor pressure at a given reference temperature^83^.

The vapor pressure is also linearly dependent on the concentration of the given compound, the partial vapor pressure follows Raoult’s law (eq. 17): the partial vapor pressure (P_i_) of each component of an ideal mixture of liquids is equal to the vapor pressure of the pure component (P_i_*) multiplied by its mole fraction in the mixture (x_i_).

Raoult’s law

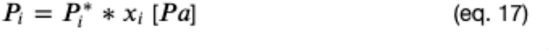

Because the boiling point is directly determined by the ambient pressure, and this pressure increases in the reactor with the depth in the broth, the pressure was expressed as a function of depth using Pascal’s law (eq. 18), assuming a broth density equal to that of water, 1000 kg/m^3^, also used to determine the pressure at the bottom of the reactor (eq. 5).

Pascal’s law

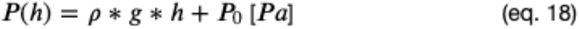

For the chosen BCR, the steady state product concentration is expressed as a function of temperature (Figure 5c). With H_2_ limited growth, ignoring product inhibition, the production rate of the compounds is related to the H_2_ uptake rate, calculated earlier, at a 9:1 H_2_ to CO_2_ ratio, using the stoichiometric yields based overall reaction balancing of CO_2_ and H_2_ as substrate and the compound of interest and water as products (Supplementary table S2). The removal rate divided by the dilution rate gives the concentration in the off-gas, which is converted to a partial pressure using the ideal gas law. From the partial pressure divided by the vapor pressure of the pure compound, a molar fraction can be calculated, assuming that a steady state is reached, which is converted to a concentration.

### 4.6 Stoichiometric model of WL pathway

All stoichiometric modeling was done using COBRApy, a package for constraints-based metabolic modeling for Python^84^. A stoichiometric model was created of the WLP with all its possible products, taking the iHN637 Genome-scale model of *Clostridium ljungdahlii* DSM 13528 as a template^85^. For investigation of downstream products of the WLP, a branching pathway to acetone production was created by adding the reactions Acetate-acetoacetate CoA-transferase (ctfAB), and acetoacetate decarboxylase (ADC), involved in acetone production by *Clostridium acetobutylicum*^86^. Sink reactions were created for all energy and redox metabolites (ATP/ADP/Pi, NADPH/NADP, NADH/NAD, Ferredoxin o/r, H^+^) (notebook S03A). Flux Balance Analysis (FBA) was used to predict fluxes. The generation and usage of energy and redox intermediates was taken directly from the predicted flux of the sink reactions (notebook 06), determined first upstream of acetyl-CoA, separately for growth on CO and H_2_, then downstream for each product. A map was created to visualize the predicted fluxes, using the Escher package (Supplementary Figure S3). This model was extended to create a second model that includes all relevant bifurcating enzymes (notebook S03B). All sinks except ATP/ADP/Pi were removed and four bifurcating enzymes were added: Ferredoxin: NADP reductase, ATP synthase, Ferredoxin NADH linked hydrogenase, and Ferredoxin:NAD oxidoreductase. For FBA, the product transportation reaction is set as the objective function and either CO_2_ or CO uptake is set to be limiting (notebook 07).

## Nomenclature

a: area [m^2^]
c: concentration (c*: saturation c, c_P_* limiting inhibitory c) [mM]
D: Diffusion coefficient [m^2^/s]
F: Flow rate [m^3^/h]
g: acceleration of gravity [m/s^2^]
H: Henry’s law constant [mol/m^3^*Pa]
h: height [m]
k: Coefficient or constant, defined per case
m_GAM_: GAM energy requirement [kJ/mol_biomass_]
m_NGAM_: NGAM energy requirement [kJ/h/mol_biomass_]
P: (vapor) pressure (P_m_: mean logarithmic, P_b_: bottom, P_t_: top, P*: partial) [Pa]
q: specific consumption or production rate [h^-1^]
R: Gas constant [m^3^*Pa/K/mol] or R Transfer rate [mol/m^3^/s]
r: Radius [m]
T: Temperature [K]
V: Volume [m^3^]
v^c^_gs_: superficial gas velocity [m^3^/h]
v: stoichiometric factor [unitless]
x: mol fraction [mol/mol]
Y: Yield (PS: product on substrate, XS: biomass on substrate, …) [(C)mol/(C)mol]
ΔH_r/f/vap_: Enthalpy of reaction/formation/vaporization [kJ/mol]
ΔG_r/f_: Gibbs free energy of reaction/formation [kJ/mol]
ε: gas holdup fraction [unitless]
θ: Temperature correction factor of k_L_a [unitless]
μ: specific growth rate [/h]
π: pi [unitless]
ρ: density [kJ/m^3^]

## Subscripts

i: compound I (or inhibition for K_i_)
X: Biomass
S: Substrate
P: Product
max: maximum
ana: anabolic reaction
cata: catabolic reaction
G: in the gas phase
L: in the liquid bulk
in: incoming (from feed flow)
T: at temperature T
0: at a specified reference condition

## Acknowledgments

We would like to thank Ivan Pogrebnyakov for fruitful discussions and support throughout the project. P.J.G.d.V. and V.M. are the recipients of a fellowship from the Novo Nordisk Foundation as part of the Copenhagen Bioscience Ph.D. Program, supported through grants NNF19CC0035454 and NNF18CC0033664 respectively. N.S. was funded by the Novo Nordisk Foundation within the framework of the Fermentation-based Biomanufacturing Initiative (FBM), grant number NNF17SA0031362. A.T.N. and T.Ø.J. received funding from the European Union’s Horizon 2020 research and innovation programme under grant agreement number 101037009 (PyroCO2). We further acknowledge funding from the Novo Nordisk Foundation (grant number NNF20CC0035580), the Villum Fonden (grant number 40986) and the Danish Research Council (grant number 103200448B).

The authors declare that there are no competing interests associated with the contents of this article.

## Author contributions

**P.J.G.d.V**.: Conceptualization, Investigation, Computational methodology, Data curation, Visualization, Writing – original draft and editing. **V.M**.: Conceptualization, Data curation, Visualization, Writing – original draft and editing. **N.S:** Writing – review, and editing. **T.Ø.J**.: Conceptualization, Supervision, Writing – review, and editing. **A.T.N**.: Conceptualization, Supervision, Funding acquisition, Project administration, Writing – review, and editing. All authors have read and approved this manuscript.

