## Supplementary Files for "Probing efficient microbial CO_2_ utilization through metabolic and process modeling"

[*] Correspondence to:

### Supplementary Figures

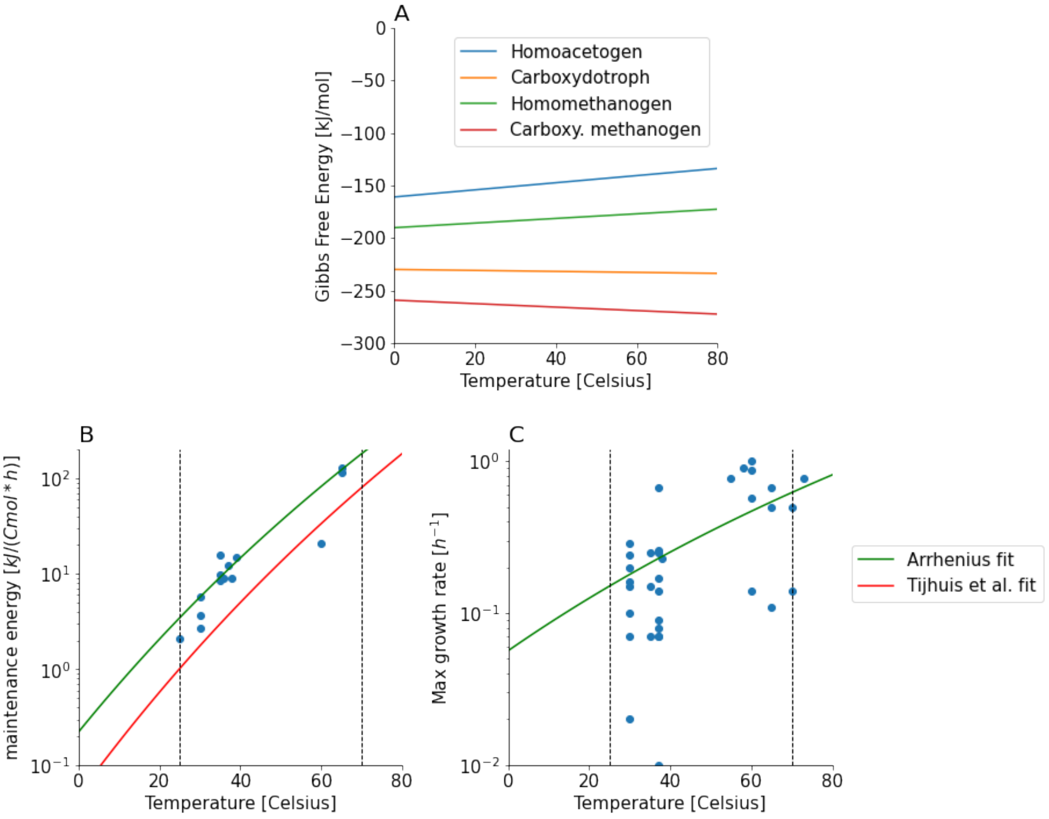


**Supplementary Figure S1:** Predicted temperature effect on (A) the $\Delta_r G$ of acetogenesis and methanogesis, (B) on the anaerobic NGAM energy requirements of anaerobic bacteria, as a function of temperature, as fitted by Thijhuis et al.[46], and (C) on the maximum growth rate, as fitted to the Arrhenius equation, with literature data represented as blue dots and listed in supplementary table 8.

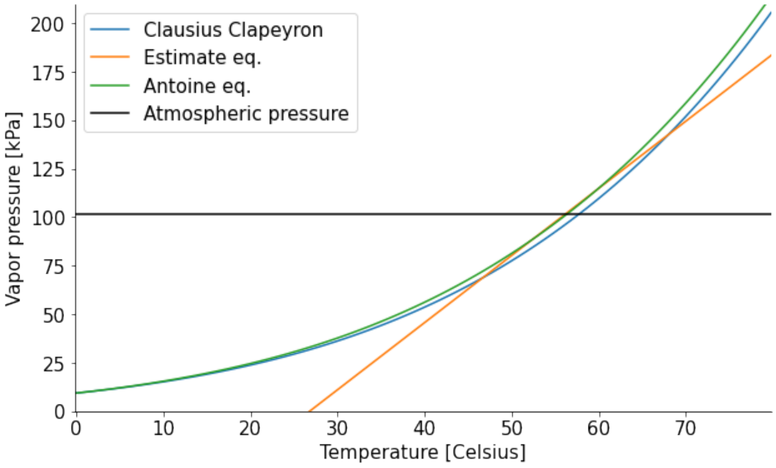


**Supplementary Figure S2:** method comparison for the temperature correction of a compound's vapor pressure, tested for acetone.

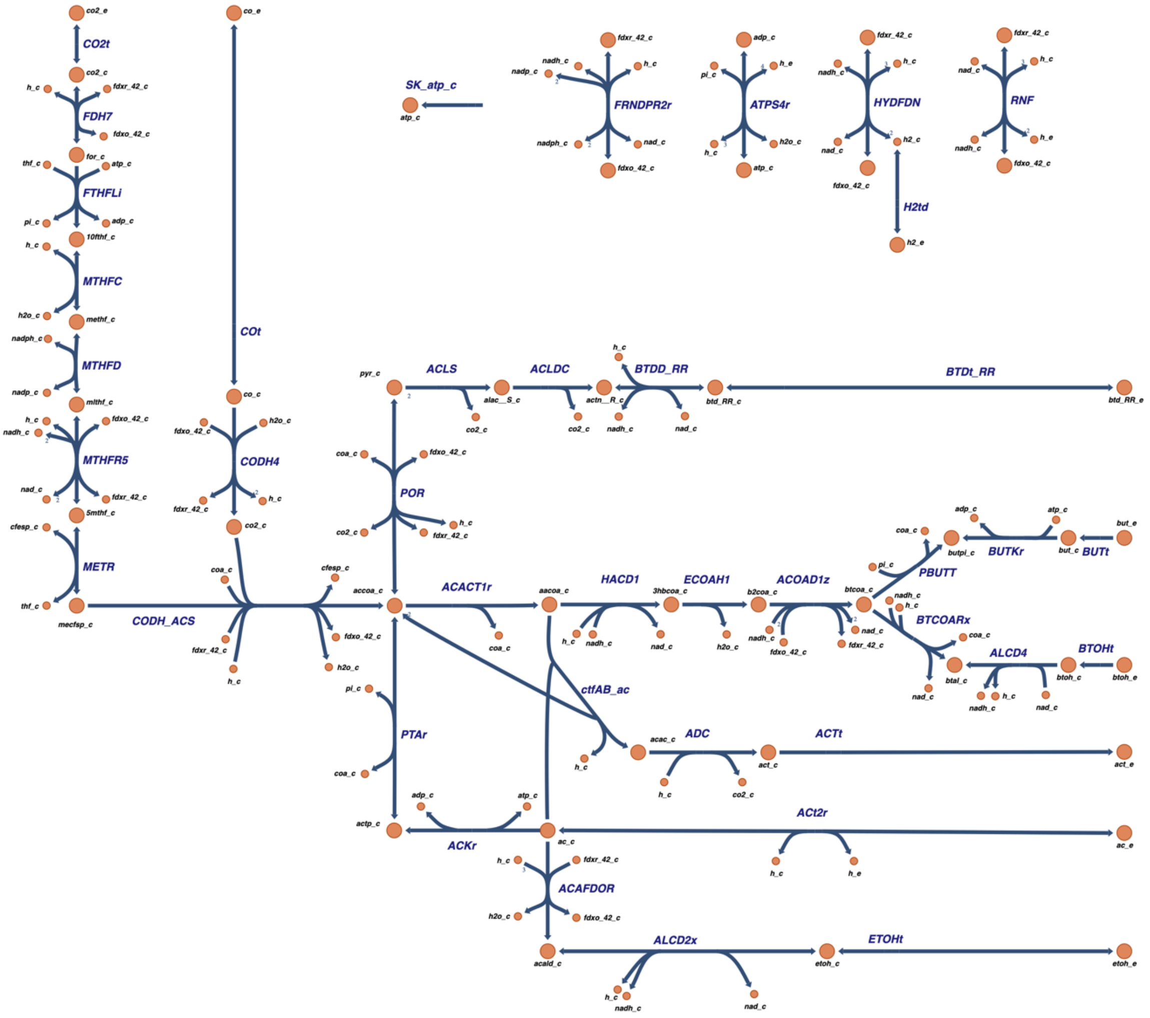


**Supplementary Figure S3:** WLP map with various possible products, including heterologously expressed acetone. Abbreviations correspond to those used in the C. ljungdahlli GEM [52]. Metabolite Abbreviations ("_c" indicates cellular and "_e" extracellular): pi: inorganic phosphate, h: H+, ac: acetate, etoh: ethanol, btd_RR: Butanediol, but: Butyrate, btoh: Butanol, fdxr_42: Ferredoxin reduced, fdxo_42: Ferredoxin oxidized, for: Formate, atp: Adenosine triphosphate (ATYP), thf: 5,6,7,8-Tetrahydrofolate, 10fthf: 10-Formyltetrahydrofolate, adp: Adenosine diphosphate (ADP), methf: 5,10-Methenyltetrahydrofolate, mlthf: 5,10-Methylenetetrahydrofolate, nadp: Nicotinamide adenine dinucleotide phosphate (NADP), nadph: Nicotinamide adenine dinucleotide phosphate

- reduced (NADPH), nadh: Nicotinamide adenine dinucleotide - reduced (NADH), 5mthf: 5-Methyltetrahydrofolate, nad: Nicotinamide adenine dinucleotide (NAD), cfesp: Corrinoid Iron sulfur protein, mecfsp: Methylcorrinoid iron sulfur protein, coa: Coenzyme A, accoa: Acetyl_CoA, actp: Acetyl P, acald: Acetaldehyde, pyr: Pyruvate, alac__S: Acetolactate, actn__R: Acetoin, aacoa: Acetoacetyl-CoA, 3hbcoa: (S)-3-Hydroxybutanoyl-CoA, b2coa: Crotonoyl-CoA, btcoa: Butanoyl-CoA, btal: Butanal, butpi: Butanoyl Phosphate, acac: Acetoacetate, act: Acetone, act: Acetone

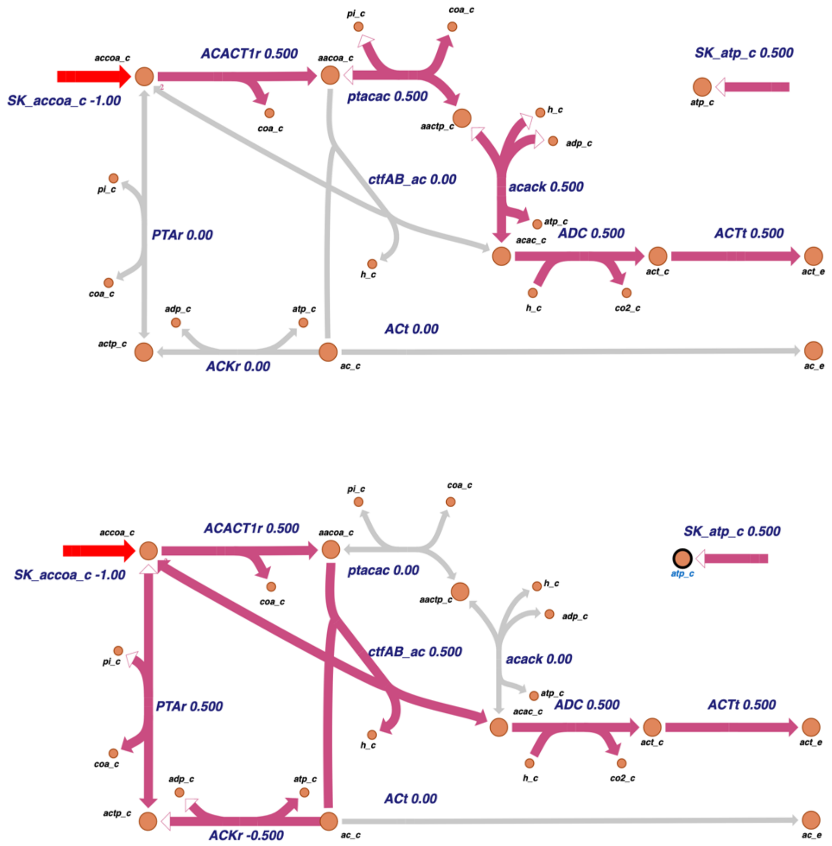


**Supplementary Figure S4:** visual comparison of two pathways from acetyl-CoA to acetone. pathway from ABE-fermentation (top) and the ATP generating pathway through acetoacetyl-P (aactp_c, bottom). ATP costs are identical, as can be read from the sink reaction in the top right corner of both maps. Metabolite abbreviations are the same as in figure S3.

#### Supplementary Tables

**Supplementary Table 1.** Reaction stoichiometry, Gibbs energy of reaction at 25°C (ΔGr), ATP yield and product yield for acetogenic formation of various products. For each product the first row is the carboxydotrophic reaction, and the second is homoacetogenic. Yields are obtained from the stoichiometric model, while the Gibbs energy of reaction is the difference in $\Delta_f G$ of products and substrates. Grey shading indicates homoacetogenesis.

| Product | Equation | ΔGr | ATP Yield | Prod. Yield |
|---|---|---|---|---|
| | | kJ/emol | Mol/emol | Cmol/emol |
| Acetate | 4 CO + 2 H2O <=> 2 CO2 + CH3COOH | -57.8 | 0.42 | 0.5 |
| | 2 CO2 + 4 H2 <=> 2 H2O + CH3COOH | -38.1 | 0.08 | 0.5 |
| Ethanol | 6 CO + 3 H2O <=> 4 CO2 + C2H5OH | -54.7 | 0.39 | 0.33 |
| | 2 CO2 + 6 H2 <=> 3 H2O + C2H5OH | -35.0 | 0.06 | 0.33 |
| Acetone | 8 CO + 3 H2O <=> 5 CO2 + C3H6O | -57.6 | 0.25 | 0.38 |
| | 3 CO2 - 8 H2 <=> 5 H2O + C3H6O | -37.9 | -0.08 | 0.38 |
| Butyrate | 12 CO + 5 H2O <=> 8 CO2 + C4H9OH | -58.5 | 0.37 | 0.4 |
| | 4 CO2 - 12 H2 <=> 7 H2O + C4H9OH | -38.8 | 0.03 | 0.4 |
| Butanol | 10 CO + 5 H2O <=> 6 CO2 + C4H8O2 | -57.1 | 0.33 | 0.33 |
| | 4 CO2 + 10 H2 <=> 6 H2O + C4H8O2 | -37.4 | 0.00 | 0.33 |
| Butanediol | 11 CO + 5 H2O <=> 7 CO2 + C4H10O2 | -49.7 | 0.12 | 0.36 |
| | 4 CO2 + 11 H2 <=> 6 H2O + C4H10O2 | -30.0 | -0.21 | 0.36 |

**Supplementary Table 2:** constants & values

| Name | Symbol = Value [unit] | Ref |
|---|---|---|
| Temperature correction factor of $k_L a$ | θ = 1.022 | [37] |
| pi | π = 3.141 | |
| Gas constant | R = 8.314 [$m^3$*Pa/K/mol] | |
| Acceleration of gravity | g = 9.81 [$m/s^2$] | |

**Supplementary Table 3:** reactor properties

| Name | Symbol = Value [unit] | Ref |
|---|---|---|
| Gas flow rate | $F_G = 1*10^4$ [$m^3/h$] | [20] |
| Reactor radius | r = 3 [m] | [20] |
| Reactor height | h = 30 [m] | [20] |
| Pressure at the top (atmospheric) | $p_t$ = 101 325 [Pa] | |

**Supplementary Table 4:** diffusion properties and Henry's law data of gaseous compounds in water

| Diffusion coefficient at infinite dilution in water at 298.15K | | REF |
|---|---|---|
| CO | $D_{0,CO} = 2.03 * 10^{-5}$ [$cm^2/s$] | 81 |
| CO2 | $D_{0,CO2} = 1.92 * 10^{-5}$ [$cm^2/s$] | 81 |
| H2 | $D_{0,H2} = 4.50 * 10^{-5}$ [$cm^2/s$] | 81 |
| O2 | $D_{0,O2} = 2.10 * 10^{-5}$ [$cm^2/s$] | 81 |
| N2 | $D_{0,N2} = 1.88 * 10^{-5}$ [$cm^2/s$] | 81 |
| Henry's law constant for solubility in water, at 298.15K | | |
| CO | $H_{0,CO} = 0.00099 * 10^3$ [$mol/m^3/Pa$] | 41 |
| CO2 | $H_{0,CO2} = 0.035 * 10^3$ [$mol/m^3/Pa$] | 41 |
| H2 | $H_{0,H2} = 0.00078 * 10^3$ [$mol/m^3/Pa$] | 41 |
| O2 | $H_{0,O2} = 0.0013 * 10^3$ [$mol/m^3/Pa$] | 41 |
| N2 | $H_{0,N2} = 0.0006 * 10^3$ [$mol/m^3/Pa$] | 41 |
| Henry's law constant temperature correction factor k | | |
| CO | $k_{CO} = 1300$ [K] | 41 |
| CO2 | $k_{CO2} = 2400$ [K] | 41 |
| H2 | $k_{H2} = 500$ [K] | 41 |
| O2 | $k_{O2} = 1500$ [K] | 41 |
| N2 | $k_{N2} = 1300$ [K] | 41 |

**Supplementary Table 5:** diffusion and phase change properties of solutes in air

| **Diffusion coefficient at infinite dilution in water at 298.15K** | | **Ref.** |
|---|---|---|
| Water | $D_{GR,H2O}$ = 0.2523 * $10^{-4}$ [$m^2$/s] | 82 |
| Acetate | $D_{GR,Actt}$ = 0.1240 * $10^{-4}$ [$m^2$/s] | 82 |
| Methanol | $D_{GR,MetOH}$ = 0.1526 * $10^{-4}$ [$m^2$/s] | 82 |
| Ethanol | $D_{GR,EtOH}$ = 0.1186 * $10^{-4}$ [$m^2$/s] | 82 |
| Propanol | $D_{GR,PropOH}$ = 0.0997 * $10^{-4}$ [$m^2$/s] | 82 |
| Butanol | $D_{GR,ButOH}$ = 0.0864 * $10^{-4}$ [$m^2$/s] | 82 |
| Butanone | $D_{GR,Butnon}$ = 0.0906 * $10^{-4}$ [$m^2$/s] | 82 |
| Butyrate | $D_{GR,Butyrt}$ = 0.0778 * $10^{-4}$ [$m^2$/s] | 82 |
| Butanediol | $D_{GR,Butdiol}$ = 0.0861 * $10^{-4}$ [$m^2$/s] | 82 |
| Acetone | $D_{GR,Actn}$ = 0.1053 * $10^{-4}$ [$m^2$/s] | 82 |
| **Enthalpy of vaporization** | | |
| Water | $\Delta_{vap}H_{H2O}$ = 4.3 * $10^{-4}$ [J/mol] | 83 |
| Acetate | $\Delta_{vap}H_{Actt}$ = 5.16 * $10^{-4}$ [J/mol] | 83 |
| Methanol | $\Delta_{vap}H_{MetOH}$ = 3.76 * $10^{-4}$ [J/mol] | 83 |
| Ethanol | $\Delta_{vap}H_{EtOH}$ = 4.2 * $10^{-4}$ [J/mol] | 83 |
| Propanol | $\Delta_{vap}H_{PropOH}$ = 4.7 * $10^{-4}$ [J/mol] | 83 |
| Butanol | $\Delta_{vap}H_{ButOH}$ = 5.2 * $10^{-4}$ [J/mol] | 83 |
| Butanone | $\Delta_{vap}H_{Butnon}$ = 3.4 * $10^{-4}$ [J/mol] | 83 |
| Butyrate | $\Delta_{vap}H_{Butyrt}$ = 5.9 * $10^{-4}$ [J/mol] | 83 |
| Butanediol | $\Delta_{vap}H_{Butdiol}$ = 7.8 * $10^{-4}$ [J/mol] | 83 |
| Acetone | $\Delta_{vap}H_{Actn}$ = 3.1 * $10^{-4}$ [J/mol] | 83 |
| **Vapor Pressure of pure compounds at 293.15K** | | |
| Water | $P_{H2O}$ = 2.4 * $10^{3}$ [Pa] | 84 |
| Acetate | $P_{Actt}$ = 1.5 * $10^{3}$ [Pa] | 85 |
| Methanol | $P_{MetOH}$ = 12.9 * $10^{3}$ [Pa] | 85 |
| Ethanol | $P_{EtOH}$ = 5.8 * $10^{3}$ [Pa] | 85 |
| Propanol | $P_{PropOH}$ = 2.0 * $10^{3}$ [Pa] | 85 |
| Butanol | $P_{ButOH}$ = 0.58 * $10^{3}$ [Pa] | 85 |
| Butanone | $P_{Butnon}$ = 10.5 * $10^{3}$ [Pa] | 85 |
| Butyrate | $P_{Butyrt}$ = 0.057 * $10^{3}$ [Pa] | 85 |
| Butanediol | $P_{Butdiol}$ = 0.010 * $10^{3}$ [Pa] | 85 |
| Acetone | $P_{Actn}$ = 24 * $10^{3}$ [Pa] | 85 |

**Supplementary Table 6:** Gibbs energy data

| **Gibbs energy of Formation and standard molar enthalpy at 298.15** | | | **Ref.** |
|---|---|---|---|
| CO2 | $\Delta_f G_{0,CO2}$ = -394.36 [kJ/mol] | $\Delta_f H_{0,CO2}$ = -393.50 [kJ/mol] | [47] |
| CO | $\Delta_f G_{0,CO}$ = -119.90 [kJ/mol] | $\Delta_f H_{0,CO}$ = -120.96 [kJ/mol] | [47] |
| H2 | $\Delta_f G_{0,H2}$ = 17.60 [kJ/mol] | $\Delta_f H_{0,H2}$ = -4.20 [kJ/mol] | [47] |
| O2 | $\Delta_f G_{0,O2}$ = 16.40 [kJ/mol] | $\Delta_f H_{0,O2}$ = 11.70 [kJ/mol] | [47] |
| Methane | $\Delta_f G_{0,Metn}$ = -34.33 [kJ/mol] | $\Delta_f H_{0,Metn}$ = -89.04 [kJ/mol] | [47] |
| Water | $\Delta_f G_{0,H2O}$ = -237.19 [kJ/mol] | $\Delta_f H_{0,H2O}$ = -285.83 [kJ/mol] | [47] |
| Acetate | $\Delta_f G_{0,Actt}$ = -396.45 [kJ/mol] | $\Delta_f H_{0,Actt}$ = -485.76 [kJ/mol] | [47] |
| Methanol | $\Delta_f G_{0,MetOH}$ = -175.31 [kJ/mol] | $\Delta_f H_{0,MetOH}$ = -23.93 [kJ/mol] | [47] |
| Ethanol | $\Delta_f G_{0,EtOH}$ = -181.64 [kJ/mol] | $\Delta_f H_{0,EtOH}$ = -288.30 [kJ/mol] | [47] |
| Propanol | $\Delta_f G_{0,PropOH}$ = -185.23 [kJ/mol] | $\Delta_f H_{0,PropOH}$ = -330.83 [kJ/mol] | [47] |
| Butanol | $\Delta_f G_{0,ButOH}$ = -171.84 [kJ/mol] | $\Delta_f H_{0,ButOH}$ = NA | [47] |
| Butyrate | $\Delta_f G_{0,Butyrt}$ = -352.63 [kJ/mol] | $\Delta_f H_{0,Butyrt}$ = NA | [47] |
| Butanediol | $\Delta_f G_{0,Butdiol}$ = -290.84 [kJ/mol] | $\Delta_f H_{0,Butdiol}$ = -505.30 [kJ/mol] | [86,87] |
| Acetone | $\Delta_f G_{0,Actn}$ = -159.70 [kJ/mol] | $\Delta_f H_{0,Actn}$ = -221.71 [kJ/mol] | [47] |

**Supplementary Table 7:** biological properties

| **Name** | **Symbol = Value [unit]** | **Ref.** |
|---|---|---|
| GAM energy requirement | $a_G$ = 1000 [kJ/CmolX] | [45] |
| Saturation constant, CO | $K_{S,CO}$ = 0.2950 [M] | est. |
| Saturation constant, H2 | $K_{S,H2}$ = 0.2950 [M] | [88] |
| Saturation constant, CO2 | $K_{S,CO2}$ = 0.2950 [M] | est. |
| Product inhibition constant, acetate for acetogens | $K_{iP,Actt}$ = 0.8 [M] (roughly 50g/l) | est. |
| max growth rate | $\mu_{max,carb}$ = 0.32 [$h^{-1}$] | [88] |

**Supplementary Table 8:** Literature values for μmax of acetogens, when not specified in the source, optimal growth temperature is taken from growth recommendation of DSMZ[57].

| **Species** | **Topt**[57] | **Reported** | **μ_max** | **Ref.** |
|---|---|---|---|---|
| *Rhodospirillum rubrum* | 30°C | 5.00 [h] | 0.20 [/h] | [89] |
| *Caldanaerobacter subterraneus* | 70°C | 7.10 [h] | 0.14 [/h] | [89] |
| *Carboxydocella sporoproducens* | 60°C | 1.00 [h] | 1.00 [/h] | [89] |
| *Carboxydocella thermoautotrophica* | 58°C | 1.10 [h] | 0.91 [/h] | [89] |
| *Carboxydothermus hydrogenoformans* | 70°C | 2.00 [h] | 0.50 [/h] | [89] |
| *Carboxydothermus islandicus* | 65°C | 2.00 [h] | 0.50 [/h] | [89] |
| *Carboxydothermus pertinax* | 65°C | 1.50 [h] | 0.67 [/h] | [89] |
| *Carboxydothermus siderophilus* | 65°C | 9.30 [h] | 0.11 [/h] | [89] |
| *Thermincola carboxydiphila* | 55°C | 1.30 [h] | 0.77 [/h] | [89] |
| *Thermolithobacter carboxydivorans* | 73°C | 1.30 [h] | 0.77 [/h] | [89] |

| | | | | |
|---|---|---|---|---|
| *Thermosinus carboxydivorans* | 60°C | 1.15 [h] | 0.87 [/h] | [89] |
| *Clostridium ljungdahlii* | 37°C | 3.8 [h] | 0.26 [/h] | [89] |
| *Clostridium autoethanogenum* | 37°C | 4 [h] | 0.25 [/h] | [89] |
| *Clostridium formicoaceticum* | 37°C | 7 [h] | 0.14 [/h] | [89] |
| *Clostridium ragsdalei* | 35°C | 4 [h] | 0.25 [/h] | [89] |
| *Clostridium scatologenes* | 37°C | 11.6 [h] | 0.09 [/h] | [89] |
| *Clostridium drakei* | 30°C | 3.5 [h] | 0.29 [/h] | [89] |
| *Clostridium carboxidivorans* | 38°C | 4.3 [h] | 0.23 [/h] | [89] |
| *Alkalibaculum bacchi* | 37°C | 5.8 [h] | 0.17 [/h] | [89] |
| *Butyribacterium methylotrophicum* | 37°C | 13.9 [h] | 0.07 [/h] | [89] |
| *Moorella thermoautotrophica* | 60°C | 7 [h] | 0.14 [/h] | [89] |
| *Oxobacter pfennigii* | 37°C | 13.9 [h] | 0.07 [/h] | [89] |
| *Acetobacterium Woodii* | 30 | 6.2 [h] | 0.16 [/h] | [89] |
| *Blautia producta* | 37 | 1.5 [h] | 0.67 [/h] | [89] |
| *Clostridium aceticum* | 30 | 10 [h] | 0.10 [/h] | [89] |
| *Acetobacterium fimetarium* | 30 | 0.45 [/d] | 0.02 [/h] | [90] |
| *Acetobacterium wieringae* | 30 | 1.71 [/d] | 0.07 [/h] | [90] |
| *Blautia hydrogenotrophica* | 37 | 1.87 [/d] | 0.08 [/h] | [90] |
| *Clostridium magnum* | 30 | 3.5 [/d] | 0.15 [/h] | [90] |
| *Eubacterium aggregans* | 37 | 0.12 [/d] | 0.01 [/h] | [90] |
| *Sporomusa acidovorans* | 35 | 1.58 [/d] | 0.07 [/h] | [90] |
| *Sporomusa ovata* | 35 | 3.59 [/d] | 0.15 [/h] | [90] |
| *Terrisporobacter mayombei* | 30 | 5.77 [/d] | 0.24 [/h] | [90] |
| *Thermoanaerobacter kivui* | 60 | 1.75 [h] | 0.57 [/h] | [91] |
